# Transcriptomic network analysis of brain and bone reveals shared molecular mechanisms underlying Alzheimer’s Disease and related dementias (ADRD) and Osteoporosis

**DOI:** 10.1101/2023.10.26.559969

**Authors:** Archana Nagarajan, Jason Laird, Obiadada Ugochukwu, Sjur Reppe, Kaare Gautvik, Ryan D. Ross, David A. Bennett, Clifford Rosen, Douglas P. Kiel, Lenora A. Higginbotham, Nicholas Seyfried, Christine W. Lary

## Abstract

Alzheimer’s disease and related dementias (ADRD) and Osteoporosis (OP) are two prevalent diseases of aging with numerous epidemiological associations, but the underlying molecular mechanisms contributing to this association are unknown. We used WGCNA (weighted gene co-expression network analysis) to develop transcriptomic networks in bone and brain tissue using two different studies to discover common molecular mechanisms. We used RNA-sequencing data from the dorsolateral prefrontal cortex tissue of autopsied brains in 629 participants from ROSMAP (Religious Orders Study and the Memory and Aging Project), including a subset of 298 meeting criteria for inclusion in five ADRD categories and the full set in a secondary analysis, and RNA array data from transiliac bone in 84 participants from the Oslo study of postmenopausal women. After developing each network, we analyzed associations between modules (groups of co-expressed genes) with multiple bone and neurological traits, examined overlap in modules between networks, and performed pathway enrichment analysis to discover conserved mechanisms. We discovered three modules in ROSMAP that showed significant associations with ADRD and bone related traits and four modules in Oslo that showed significant associations with multiple bone outcomes. We found significant module overlap between the two networks, most notably among those modules linked to canonical Wnt signaling and skeletal tissue homeostasis and development. These results were preserved with a network from the full ROSMAP cohort (n=629), which included a broader spectrum of participants. Our results require validation in experimental studies but show support for Wnt signaling as an important driver of pathology in OP and ADRD. We additionally show a strong link between Dementia with Lewy bodies and bone outcomes. These results have translational significance in the development of novel treatments and biomarkers for both ADRD and OP.

## Introduction

Alzheimer’s disease and related dementias (ADRD) represent a group of progressive age-related neurodegenerative disorders that are typically characterized by memory loss, cognitive decline, and impaired daily functioning. Alzheimer’s disease (AD) accounts for approximately 60-80% of all cases of ADRD, making it the most common type of dementia, above Lewy Body dementia, vascular dementia, and frontotemporal dementia [1, 2]. Patients with ADRD have shown high rates of osteoporosis (OP), a skeletal disorder characterized by reduced bone mineral density (BMD) and increased susceptibility to fractures [3, 4]. Lower BMD and increased fracture rate, especially in the hip, wrist, and spine, have been reported in ADRD patients [5, 6].

On a molecular level, several mechanisms have been shown to contribute to the pathophysiology of OP and ADRD. These mechanisms include oxidative stress [7], accumulation of advanced glycation end products (AGEs) [8], calcification [9], calcium ion imbalance [10], immunological factors such as cytokine secretion [11–13], defects in glucose uptake [14], endoplasmic reticulum-stress driven senescence [15], endogenous or exogenous estrogen exposure [16–19], vitamin D signaling [20, 21], receptor activator of nuclear factor-κB (RANK), RANK ligand (RANKL) and osteoprotegerin (OPG) signaling [22], and Wnt/β-catenin signaling [23–25]. Bone metabolic markers have shown association with AD outcomes, including serum osteocalcin, osteopontin, and sclerostin, and urine deoxypyridinoline/creatinine ratio and calcium/creatinine ratio [26–30]. Additionally, the genes *APP* and *BACE1* [31], *TREM2* [32–36], *FNDC5* (Irisin) [37–42], and *SNCA* (alpha synuclein) [43–47] have shown associations with both OP and ADRD through changes in gene expression. Despite several molecular mechanisms that could contribute to the co-occurrence of OP and ADRD, the relative contributions of each pathway are not known.

We used transcriptomic data from brain and bone tissue from two independent epidemiological studies (The Religious Order Study and Memory and Aging Project or “ROSMAP”, and The Oslo Study of Post-Menopausal Women or “Oslo”) to discover common mechanisms that may contribute to ADRD pathogenesis and osteoporotic bone outcomes [48, 49]. The use of gene co-expression networks, such as those created by weighted gene co-expression network analysis (WGCNA), allow us to identify and study groups of co-expressed genes that correlate with a specific trait [50]. We hypothesize that WGCNA of both studies, ROSMAP and Oslo, will reveal common mechanisms and gene interactors that contribute to both ADRD and OP. Undercovering shared mechanisms can lead to the discovery of novel therapeutics and biomarkers for each disease.

## Results

### ROSMAP and Oslo Cohort descriptions

The cohort characteristics are summarized in **Table 1**. 629 ROSMAP participants met our inclusion criteria with a median age of 89, 64% female. We specifically defined five categories based on the pathological criteria for Alzheimer’s disease (AD) or for Dementia with Lewy Bodies (DLB) with neuropathological changes (NC) and who were either symptomatic or asymptomatic, or controls without cognitive impairment (see **Table S1** and Methods for details). This resulted in a subgroup of 298 participants with similar characteristics (median age 89, 63% female), which was used for our primary ROSMAP network construction. The median age of the Oslo cohort was 62, 33% of whom met the diagnostic criteria for OP [51].

**Table 1.**
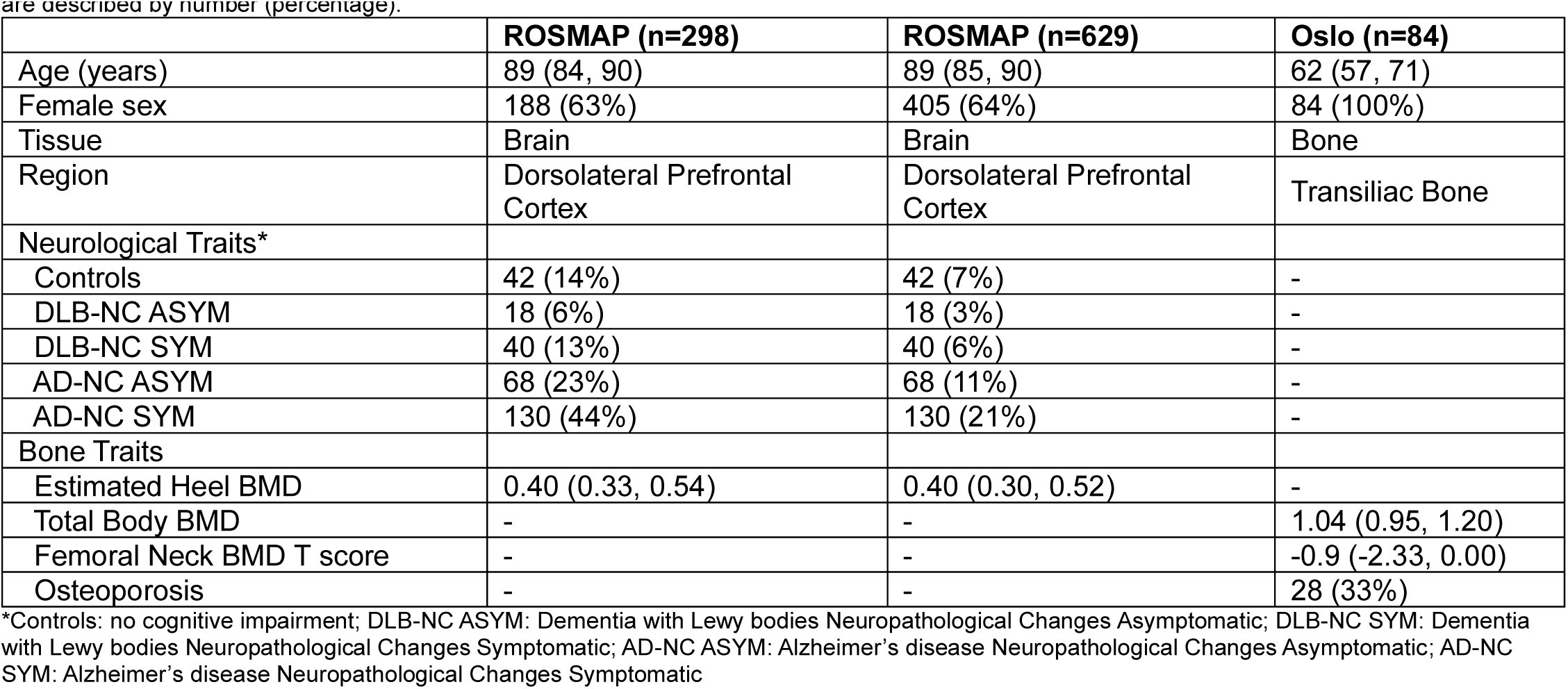
ROSMAP and Oslo Study Cohort Descriptions. Continuous variables are described by median (interquartile range) and categorical variables are described by number (percentage).

|  | **ROSMAP (n=298)** | **ROSMAP (n=629)** | **Oslo (n=84)** |
| --- | --- | --- | --- |
| Age (years) | 89 (84, 90) | 89 (85, 90) | 62 (57, 71) |
| Female sex | 188 (63%) | 405 (64%) | 84 (100%) |
| Tissue | Brain | Brain | Bone |
| Region | Dorsolateral Prefrontal Cortex | Dorsolateral Prefrontal Cortex | Transiliac Bone |
| Neurological Traits* |  |  |  |
| Controls | 42 (14%) | 42 (7%) | - |
| DLB-NC ASYM | 18 (6%) | 18 (3%) | - |
| DLB-NC SYM | 40 (13%) | 40 (6%) | - |
| AD-NC ASYM | 68 (23%) | 68 (11%) | - |
| AD-NC SYM | 130 (44%) | 130 (21%) | - |
| Bone Traits |  |  |  |
| Estimated Heel BMD | 0.40 (0.33, 0.54) | 0.40 (0.30, 0.52) | - |
| Total Body BMD | - | - | 1.04 (0.95, 1.20) |
| Femoral Neck BMD T score | - | - | -0.9 (-2.33, 0.00) |
| Osteoporosis | - | - | 28 (33%) |

\*Controls: no cognitive impairment; DLB-NC ASYM: Dementia with Lewy bodies Neuropathological Changes Asymptomatic; DLB-NC SYM: Dementia with Lewy bodies Neuropathological Changes Symptomatic; AD-NC ASYM: Alzheimer's disease Neuropathological Changes Asymptomatic; AD-NC SYM: Alzheimer's disease Neuropathological Changes Symptomatic

### Discovered WGCNA networks in bone and brain tissue

Following preprocessing of the transcriptomic data (**Figure 1A**), WGCNA of the ROSMAP subgroup (298 samples) yielded a network comprising 13,975 genes organized into 28 modules (**Figure 1B, 1C**, **Table S2**). When applied separately to the Oslo cohort, WGCNA generated a network comprising 10,381 genes and 22 modules (**Figure 1D, 1E**, **Table S3**). The network for the full ROSMAP cohort with all 629 samples had 13,975 genes with 23 modules **(Figure S1, Table S4).** Modules of co-expressed genes were labeled with a color designation as part of the WGCNA pipeline.

**Figure 1.**
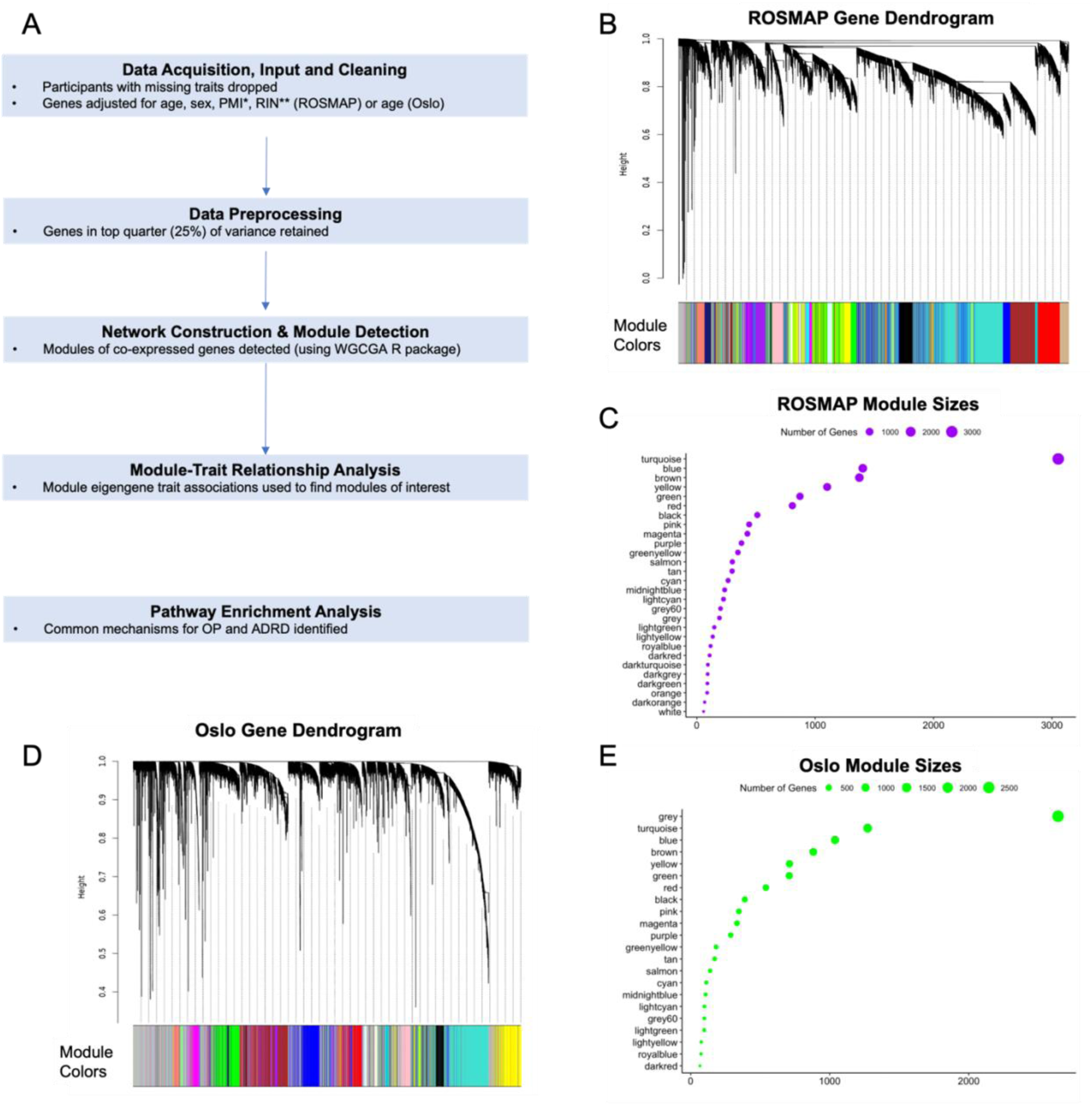
WGCNA pipeline and network construction. (A) Schematic overview of WGCNA workflow. (B) Gene dendrogram of ROSMAP network, where colors represent the labels of modules generated in the network. (C) Plot of number of genes that correspond to each module in ROSMAP’s network. (D) Gene dendrogram of Oslo network. (E) Plot of number of genes that correspond to each module in Oslo’s network.

### Module-trait analysis reveals modules correlated with neurological and bone traits

In the ROSMAP subgroup, modules with eigengenes that were significantly correlated with Heel BMD and two ADRD traits were selected for analysis. The red module (807 genes) was inversely correlated with heel BMD (r=-0.17, p=0.004) and positively correlated with multiple ADRD traits (AD-NC SYM, r=0.12, p=0.03; DLB-NC ASYM, r=0.2, p=8e-07; DLB, r= 0.14, p=0.014; amyloid, r=0.23, p=8e-05) with a trend towards significant correlation with DLB-NC SYM (r=0.11, p=0.06). The cyan module (264 genes) was positively correlated with heel BMD (r=0.16, p=0.006) and inversely correlated with ADRD traits (DLB-NC ASYM r=-0.16, p=0.005; DLB, r=-0.13, p=0.003; amyloid, r=-0.18, p=0.002), with a trend towards a significant inverse correlation with DLB-NC SYM (r=-0.11 p=0.06) (**Figure 2A**). The light yellow module (264 genes) was positively correlated with heel BMD (r=0.12, p=0.04), inversely correlated with ADRD traits (DLB-NC SYM, r=-0.18, p=0.002; DLB, r=-0.15, p=8.7e-03; amyloid, r=-0.14, p=0.02), and showed a trend towards significant inverse correlation with AD-NC ASYM (r=-0.11 p=0.07). Overall, it the modules of interests (significant for bone and ADRD traits) showed more significant correlations with DLB-NC traits compared with AD-NC traits.

**Figure 2.**
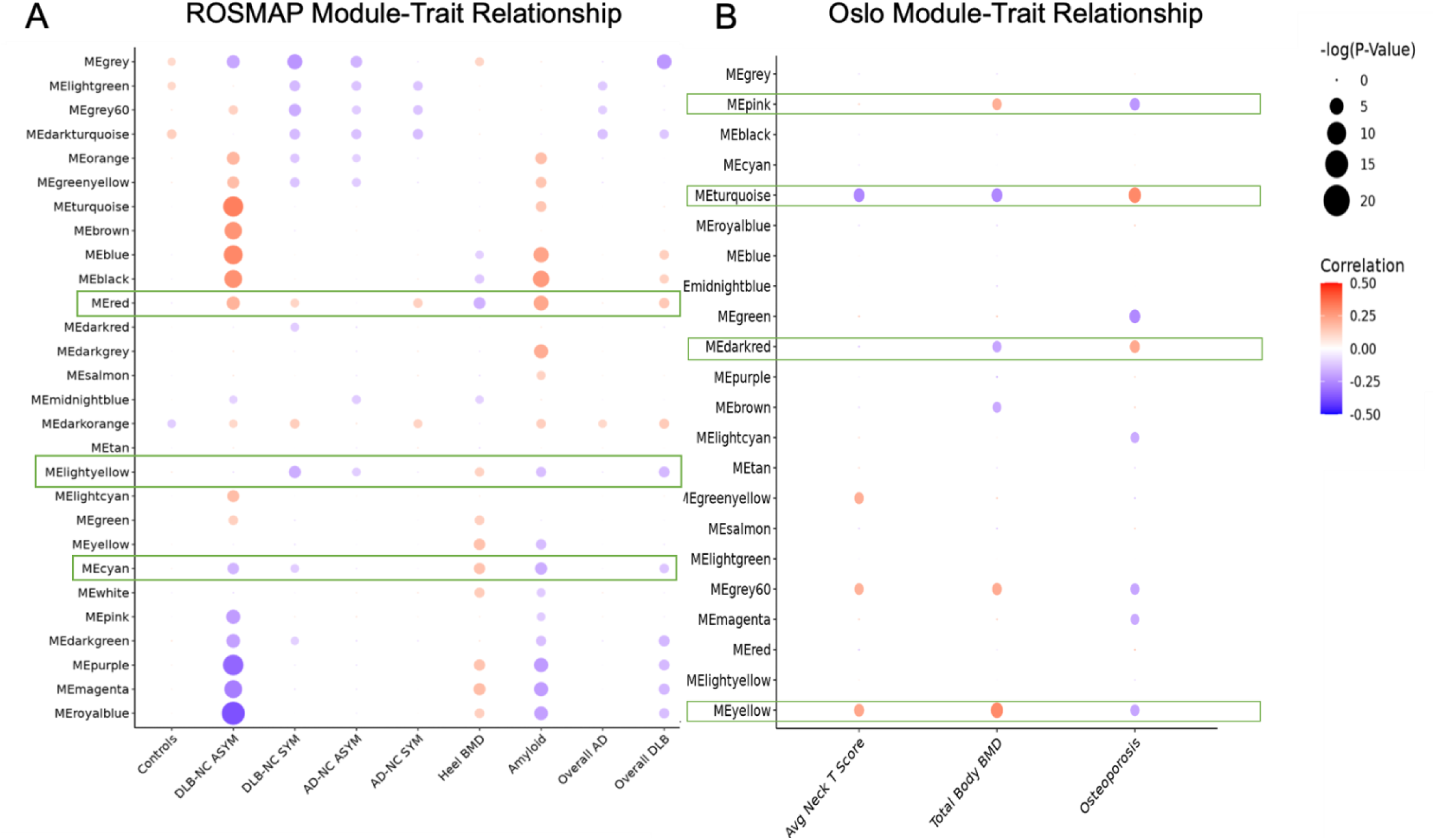
Module-trait analysis of ROSMAP and Oslo networks reveal modules of interest. (A) Dotplot of module eigengene correlation with ADRD and bone traits in ROSMAP network. Correlation values are noted through dot color and *p* value through dot size. Values shown are for -log(*p),* where p< 0.1. (B) Dotplot of module eigengene correlation with osteoporotic and bone traits in Oslo network. Red boxes represent modules of interest.

In Oslo the turquoise module (1,273 genes) was positively correlated with OP (r=0.30, p=0.005) and inversely correlated with total body BMD (r=-0.26, p=0.02) and femoral neck T score (r=-0.26, p=0.02) (**Figure 2B**). The pink module (347 genes) was inversely correlated with OP (r=-0.22, p=0.04) and showed a trend toward significant correlation with total body BMD (r=0.21, p=0.06) (**Figure 2B**). The yellow module (711 genes) showed a trend towards significant inverse correlation with OP (r=-0.19, p=0.08) and was positively correlated with total body BMD (r=0.30, p=0.006) and femoral neck T score (r=0.23, p=0.04) (**Figure 2B**). The dark red module (67 genes) was positively correlated with OP (r=0.22, p=0.04) and showed a trend towards significant inverse correlation with total body BMD (r=-0.19, p=0.08) (**Figure 2B**).

We discovered modules that were enriched for a wide array of molecular functions (**Table S5**, **Table S6** for ROSMAP and Oslo, respectively). The resulting significant GO terms for each module were manually annotated and grouped into major categories (**Figure 3**). The ROSMAP network modules grouped into basic cellular categories (DNA/RNA Metabolism, Immune Processes) and categories specific to brain tissue (Synapse Homeostasis, Neural Cell Development and Homeostasis) (**Figure 3A**). The Oslo network modules grouped into basic categories with annotations like those of ROSMAP (Immune Processes, Protein Regulation) and into a category specific to bone tissue (Skeletal Development & Signaling Pathways) (**Figure 3B**).

**Figure 3.**
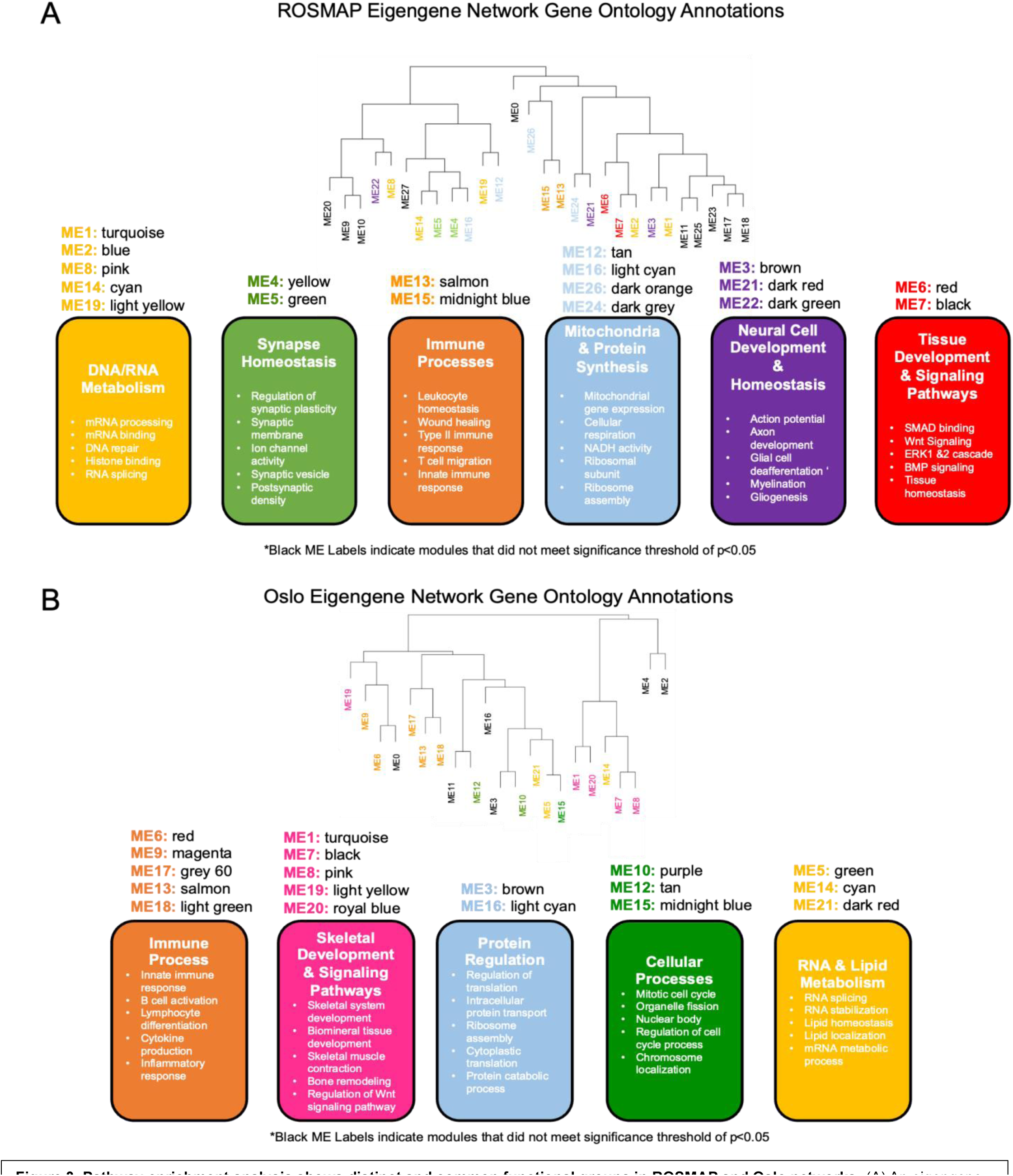
Pathway enrichment analysis shows distinct and common functional groups in ROSMAP and Oslo networks. (A) An eigengene network was constructed by hierarchical clustering of 28 modules from ROSMAP study. Modules are grouped into functional categories based on GO term enrichment. (B) Eigengene network of 22 modules from Oslo study. Modules are grouped into functional categories based on GO term enrichment.

### Overlap in modules between networks

The red module from the ROSMAP subgroup network had significant overlap with three of the four modules of interest in the Oslo network (pink, turquoise, and yellow), and the ROSMAP cyan module had significant overlap with the yellow module from Oslo. We then selected the ROSMAP red and cyan modules and the Oslo pink, turquoise, and yellow modules for further study as these modules showed associations for selected bone and neurological traits and showed significant overlap across networks (**Table S7**, **Table S8**, **Figure S2**). Test for significance of the overlap between modules of different networks can be described in detail in the Methods section.

### Pathway analysis reveals enrichment for skeletal/tissue development and signaling pathways in modules of interest

We then examined the enriched GO terms for the categories containing our modules of interest. The red module clustered into the “Tissue Development & Signaling Pathways” category, which included modules showing enrichment for Wnt Signaling, tissue homeostasis, and tissue remodeling, whereas the cyan module clustered into the “DNA/RNA Metabolism” category, which showed enrichment for histone binding, RNA splicing, and mRNA processing (**Figure 3A**). The pink and turquoise modules of the Oslo network clustered into the “Skeletal Development & Signaling Pathways”, which showed enrichment for skeletal system development, bone remodeling, and regulation of Wnt signaling pathway, yielding a similar profile to the ROSMAP network’s red module. The yellow module from the Oslo network showed no significant enrichment of GO terms (**Figure 3B**). The GO pathway enrichment analysis between modules revealed that ROSMAP’s red module showed strong overlap with Oslo’s pink and turquoise modules. A Venn diagram was constructed to identify the common GO terms between the modules previously described from the ROSMAP and Oslo networks (**Figure 4A**, **4B**). ROSMAP red (701 terms) and Oslo pink (423 terms) showed the greatest overlap of 252 terms (**Figure 4A**). ROSMAP red (701 terms) and Oslo turquoise (262 terms) had a modest overlap of 28 terms (**Figure 4B**), confirming that ROSMAP’s red module and Oslo’s pink module showed the greatest correspondence between networks. The enriched GO terms for both the ROSMAP red and Oslo pink modules showed significant enrichment for Wnt signaling terms (**Figure 4C**, **Figure 4D**). The module overlap between networks is summarized in **Table 2**. Overall, our analysis revealed two specific modules of interest from the ROSMAP and Oslo networks, red and pink respectively, to further study.

**Figure 4.**
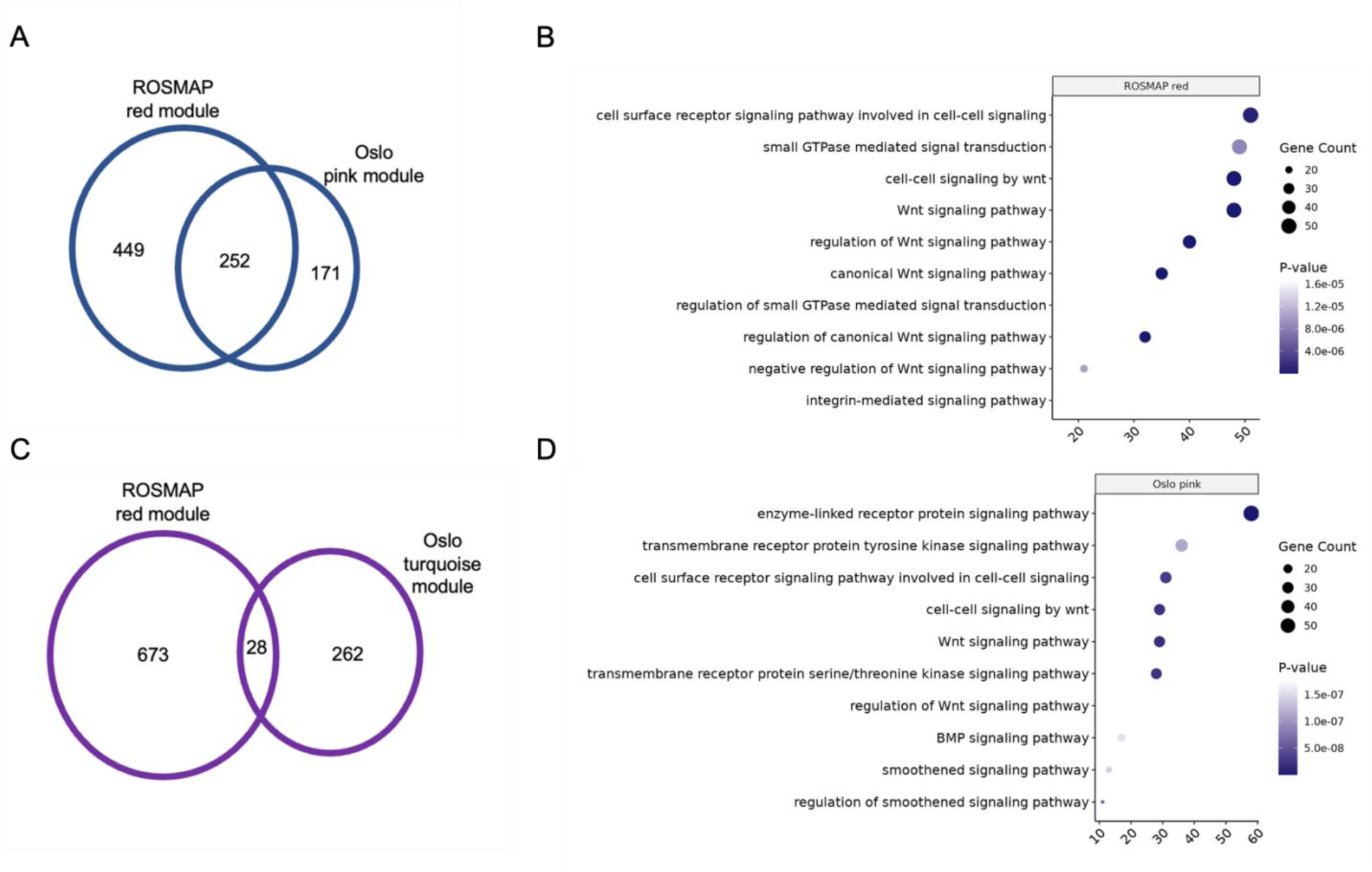
ROSMAP red and Oslo pink modules show correspondence in enriched pathways with a specific enrichment in Wnt signaling. (A) Venn diagram of ROSMAP red module and Oslo pink module enriched GO term overlap. (B) Venn diagram of ROSMAP red module and Oslo turquoise module enriched GO term overlap. (C) Dotplot of top 15 most significant (*p* value) “signal” GO terms in ROSMAP red module arranged by *p* value and gene count. (D) Dotplot of top 15 most significant (*p* value) “signal” GO terms in Oslo pink module arranged by *p* value and gene count.

**Table 2.** Overlapping modules from ROSMAP and Oslo networks. Modules showing significant overlap between networks are listed along with pathway analysis annotations. Bold-faced modules also show overlap in annotations.

| Network 1 | Module 1 | Module 1 Category | Network 2 | Module 2 | Module 2 Category |
| --- | --- | --- | --- | --- | --- |
| ROSMAP (n=298) | **red** | Tissue Development & Signaling Pathways | Oslo (n=84) | **pink** | Skeletal Development & Signaling Pathways |
|  |  |  |  | turquoise | Skeletal Development & Signaling Pathways |
|  | cyan | DNA/RNA Metabolism |  | yellow | none |
| ROSMAP (n=298) | **red** | Tissue Development & Signaling Pathways | ROSMAP (n=629) | **black** | Tissue Development & Signaling Pathways |
|  | cyan | DNA/RNA Metabolism |  | purple | none |
|  |  |  |  | grey60 | none |
| ROSMAP (n=629) | **black** | Tissue Development & Signaling Pathways | Oslo (n=84) | **pink** | Skeletal Development & Signaling Pathways |
|  |  |  |  | turquoise | Skeletal Development & Signaling Pathways |
|  | purple | none |  | yellow | none |

### Additional analysis in full ROSMAP cohort reveals consistency with subgroup results

In the ROSMAP network with the full set of participants, the black module (670 genes) was inversely correlated with heel BMD (r=-0.12, p=0.003) and positively correlated with multiple neurological traits (AD-NC SYM, r=0.1, p=0.009; DLB-NC ASYM, r=0.23, p=1e-16; DLB, r= 0.14, p=6e-04; amyloid, r=0.24, p=1e-09; DLB-NC SYM, r=0.083, p=0.04). The purple module (264 genes) was positively correlated with heel BMD (r=0.13, p=0.001) and inversely correlated with DLB-NC ASYM (r=-0.18, p=4e-06), DLB-NC SYM (r=-0.12, p=0.003), overall DLB (r=-0.14, p=4e-04), and amyloid (r=-0.18, p=4e-06) (**Figure S3**). The black module had significant overlap with the Oslo pink, turquoise, and yellow modules, and the purple module had significant overlap with the yellow module from Oslo (**Table S9**, **Table S10**, **Figure S4**), yielding similar results to the ROSMAP (subgroup) network’s red and cyan modules, which overlapped with the same set of Oslo modules, respectively. The full ROSMAP network black module overlapped with the subgroup ROSMAP network red module (**Table S11, Table S12, Figure S5)**, and the black module had similar GO term enrichments (**Table S13**) as the red module (**Table S7**), revealing consistency in our findings between the two ROSMAP networks.

### Common genes in ROSMAP, Oslo modules show links to ADRD, OP, and Wnt signaling

The ROSMAP red and Oslo pink modules fell into pathway analysis categories that were enriched for “signaling pathways”. GO pathways containing “signal” were sorted by lowest p value and highest gene count, and the top 15 pathways included several Wnt pathway GO terms (**Figure 4C**, **Figure 4D** for ROSMAP and Oslo, respectively). ROSMAP red and Oslo pink modules had 49 gene members (p=6.5e-15) that were shared (**Table S5**). Several of these genes have established associations with ADRD pathophysiology and bone outcomes (**Table S14**). Several genes were found in the Wnt pathway (*APCDD1, BAMBI, FZD8, GL13, LRP4, TSPAN6, WIFI1, ZNRF3*), which showed significance in our pathway analysis for both modules (**Figure 3**) and is further shown when genes associated with any GO term that included “Wnt” were plotted for the ROSMAP red and Oslo pink modules **(Figure S6A, Figure S6B)**. Additionally, these 49 genes have broad functional ties to tissue development, tissue morphogenesis and a wide spectrum of signaling pathways.

## Discussion

In this study we developed and compared transcriptomic networks from brain and bone tissue from ROSMAP and Oslo studies using WGCNA, in which we generated modules of co-expressed genes to discover genes and pathways associated with ADRD and OP. We discovered three modules in ROSMAP that show associations with ADRD and bone related traits, and four modules in Oslo that showed significant associations with multiple bone outcomes. We then found significant overlap between modules across the two networks, and further filtered to the overlapping modules. We performed pathway analysis of each of these modules and then grouped these into categories with related GO terms. Our major result is that the ROSMAP red module overlapped mostly strongly with the Oslo pink module both at the individual gene and GO term level, and the modules showed an enrichment for skeletal and general tissue homeostasis and development with canonical Wnt signaling pathway as a common denominator. Importantly, these relationships were preserved during our additional analysis using the unselected full group of ROSMAP participants, which included those that did not meet one of our ADRD criteria.

Interestingly, module-trait analysis showed our modules of interest (red [Tissue Development and signaling], lightyellow, cyan [DNA/RNA Metabolism]) in ROSMAP show stronger associations with DLB compared to AD neuropathological changes. All ADRD traits showed inverse correlation with heel BMD, and the correlation was strongest in the DLB-NC ASYM group - an unexpected result (**Figure 2A**). In a longitudinal follow-up study from the Korean National Health Insurance Service-Health Screening Cohort database from 2002-2015 with 78,994 OP participants and 78,994 controls, OP had a 1.27-fold higher occurrence of AD and a 1.49-fold higher occurrence of PD, indicating a slightly larger risk for developing PD [52], which involves Lewy body pathology. Interestingly, the DLB-NC ASYM trait, consisting of participants without cognitive impairment but with corticolimbic Lewy body pathology, showed the strongest inverse correlation with heel BMD. It is well established that the pathologies of neurodegenerative disease often occur years prior to the onset of cognitive impairment, indicating that bone biomarkers may be useful to screen for DLB [53, 54]. We should note that one of these groups (DLB-NC ASYM) only had a sample size of 18 limiting the generality of these results. Current research remains sparse on the bone effects of DLB compared with AD, and our study highlights the potential diversity of bone related effects of ADRD.

Amyloid beta levels were included in our study due to evidence of amyloid beta contributing to bone outcomes. A study by Li et al showed that Aβ42 and amyloid precursor protein (APP) levels were elevated in osteoporotic bone tissue from human and ovariectomized rats when compared to age and sex matched controls [55].

Another study paradoxically showed a protective effect of amyloid beta, where it promoted bone formation in ovariectomized mice by regulating Wnt/β-catenin signaling and the OPG/RANKL/RANK system [56]. Our results from the module-trait analysis in ROSMAP revealed a strong inverse correlation of heel BMD with brain amyloid beta levels. Importantly, modules that were associated with lower levels of BMD were also associated with higher levels of amyloid beta in brain tissue, indicating a negative association of amyloid beta in the brain with bone health. This study showed a strong inverse correlation between amyloid and BMD underscoring the importance of analyzing amyloid and its effects on skeletal health.

The gene overlap between the ROSMAP red and Oslo pink modules revealed several genes that have documented brain and bone effects, including *A2M, DDR2, LRP4, SOX9,* in addition to several Wnt pathway associated genes: *APCDD1*, *BAMBI*, *FZD8*, *GL13*, *LRP4*, *TSPAN6*, *WIFI1*, *SOX9*, *ZNRF3* (**Figure S6A, Figure S6B, Table S14**). A recent study showed evidence of alterations in the canonical Wnt/B catenin pathway in htau mice, a mouse line able to express six isoforms of human tau and not murine tau, compared to C57BL/6J controls [24]. The study showed reductions in Wnt receptors (LRP5/6) in brain and bone tissue with Wnt antagonists (DKK1/SOST) in bone [24]. Changes in the Wnt/β-catenin have been implicated in both ADRD and OP separately [57–61]. In comparison, our study documented this pathway to affect both brain and bone phenotypes concurrently. In a study of community-dwelling older adults with cognitive deficits, circulating levels of DKK1, a Wnt signaling pathway inhibitor, were found to be positively related to semantic memory, but negatively to working memory [61]. Although the genes of interest from these studies were not found in the genes that overlapped between the ROSMAP red and Oslo pink modules, *DKK1* and *SOST* were present in the Oslo pink module. In ROSMAP, *DKK1* was present in the turquoise module, which did not meet our module of interest criteria, and *SOST* did not make the gene list for WGCNA network construction after variance filtering. Our analysis shows a strong enrichment for Wnt signaling, particularly with the canonical Wnt/ β-catenin pathway in ADRD/OP associated modules, in concurrence with these prior studies. Strikingly, these results were found in both brain (DLPFC) and bone (transiliac) tissue and across two independent human cohort studies, increasing the strength of this finding. To our knowledge, this is the first study in humans in which the Wnt signaling pathway has been shown to be of importance in both ADRD and OP outcomes, in agreement with previous observations in animal models [24].

Interestingly, *LRP4*, *WIF1,* and *SOX9* are present in the ROSMAP and Oslo modules of interest, as each are imperative in the canonical and noncanonical Wnt pathways with documented effects in both ADRD and OP. *LRP4* was reportedly reduced in post-mortem AD brains, its genetic deletion in AD mice exacerbated cognitive deficits and increased amyloid aggregates, and its presence increased astrocytic amyloid beta uptake [62].

*LRP4* has also been seen as a therapeutic candidate for increasing bone mass, as mice with LRP4 overexpression exhibited increased bone strength and formation [63]. SOX9 has been reported to be increased in post-mortem AD brains, and one study showed improved cognition in AD mice in an miR-222-3p dependent manner through its targeting of SOX9 [64, 65]. SOX9 has also been implicated as a biomarker for primary bone cancers, and its expression was reported to be essential during musculoskeletal system development [66, 67]. Taken together, the Wnt-proteins LRP4 and SOX9 are viable candidates for further study to decipher the brain/bone connection, which may elucidate how the Wnt signaling pathway influences the pathogenesis of both ADRD and osteoporosis.

There were several other genes with brain/bone connections that were present in our ROSMAP and Oslo networks, including *FNDC5, TREM2, OPG* (osteoprotegerin), and *BGLAP* (osteocalcin) [4, 32, 35, 37, 38, 42, 68–70]. None of these genes appeared in our ROSMAP modules of interest, however *BGLAP* (osteocalcin) was present in the Oslo pink module, our main module of interest, and *FNDC5* was a member of the turquoise Oslo module, a module that met our module of interest criteria.

*APOE*, the most prominent and well-established genetic risk factor for AD representing half of AD cases [71–73], was not a member of any modules of interest. Studies have shown that the APOE ε4 allele to be associated with lower bone formation and increased risk of osteoporosis and bone fractures [74]. Additionally, SNCA, the protein responsible for Lewy Body aggregates in DLB [75], was not present in any of our modules of interest. Though the associations between SNCA and bone are not well established, one study observed that ovariectomized mice deficient in SNCA had a 40% reduction in bone loss compared to controls, and *SNCA* was found to be a central gene in the regulation of bone homeostasis and bone loss from ovariectomy [46]. Although these two key genes for AD and DLB, respectively, were not present in our modules of interest for ADRD and bone related outcomes, our work highlights novel gene targets and the importance of Wnt signaling for further study.

The joint analysis of ROSMAP and Oslo studies offers a unique and innovative approach to study the common mechanisms of ADRD and OP that have historically been challenging given the lack of detailed ADRD measures in osteoporotic cohorts, and the lack of bone measures in ADRD cohorts. Given the availability of heel BMD in participants from ROSMAP, we were able to use both bone and neurological traits in our module-trait analysis. By requiring the module eigengene to be associated with heel BMD and two other ADRD outcomes in ROSMAP and finding an OP associated module from the Oslo study that overlapped, we discovered genes that may be important for bone effects in ADRD. Additionally, this provides a unique way to add neurological context to OP cohorts, where neurological traits are typically not measured.

Our study has several weaknesses that should be noted. First, bone and brain traits were measured in different individuals in different studies and using different sequencing platforms (RNA sequencing vs RNA array). In addition, the sample sizes of individuals with OP or ADRD were relatively small. Our results are purely correlative, giving the ability to measure associations and not causal mechanisms. Finally, our study does not have experimental verification of key findings, although this will be a vital next step. The microarray analyses may not detect rare mRNAs, and the present results of co-expressed genes between the brain and bone should therefore be considered as a minimum. Our study also has strengths involving two high quality epidemiological studies of ADRD and bone outcomes in well characterized cohorts allowing for detailed assessment of module trait associations. Also, the presence of detailed pathological, cognitive, and molecular data in the ROSMAP study allows for richer interpretation of gene network results.

In summary, our study reveals an important overlap in molecular pathways underlying bone and ADRD outcomes, with a notably strong relationship between Dementia with Lewy Bodies and amyloid beta levels in the brain with BMD. This is the first human study that shows genes belonging to the Wnt pathway, of essential importance in brain and bone functions, to have strong associations with both OP and ADRD. The results warrant further study of *LRP4* and *SOX9*, two genes associated with Wnt signaling shared in our modules of interest in ROSMAP and Oslo, since both have well-documented significance to both brain and bone phenotypes. Further studies are needed to validate these associations in *in vivo* and *in vitro* models. Deciphering the crosstalk between brain and bone outcomes could lead to the discovery of important biomarkers for ADRD disease prevention or the development of new therapeutics for both OP and ADRD.

## Methods

### Cohort overview

*The Rush Alzheimer’s Disease Center* has initiated several longitudinal mixed-sex cohort studies including the Religious Orders Study (ROS) in 1994 and the Memory and Aging Project (MAP) in 1997, known jointly as ROSMAP. Each cohort recruits older participants without known dementia who sign an Anatomic Gift Act for organ donation. They include more than 3,700 participants with a follow-up rate of >90% and an autopsy rate of >85% with more than 1,800 autopsies [48] (**Table 1**). *The Oslo Study of Post-Menopausal Women* is a cohort of Norwegian women between the ages of 50-86 [49, 76]. Healthy participants and those with postmenopausal osteoporosis were recruited through the Lovisenberg Diakonale Hospital outpatient clinic as described in [76].

### Transcriptomic data

In ROSMAP, mRNA expression of dorsolateral prefrontal cortex tissue of autopsied brains was performed with RNA sequencing using the Broad Institutes’ Genomics Platform at the New York Genome Center with Illumina HiSeq, and more recently with NovaSeq in the RADC Sequencing Core [77]. The initial data processing steps are described in De Jager et al [77], with RNA integrity scores (RIN) ≥ 5 required. In Oslo, mRNA from transiliac bone biopsies was assayed using the Affymetrix HG-U133 Plus 2.0 array [49]. These samples are a heterogeneous cell population containing, in addition to bone cells, marrow cells of the haematopoietic and immunological lineages. The initial data preprocessing steps are described in Reppe at al [76], with RNA integrity scores (RIN) ≥ 5 required.

### Trait descriptions and ADRD categories

In ROSMAP, data was acquired from autopsied brain of post-mortem participants with several pathological variables including CERAD (Consortium to Establish a Registry for Alzheimer’s Disease for Neuritic Plaque Measures) score and BRAAK staging (system for measuring neurofibrillary tangle spread) available to guide diagnosis of ADRD [78, 79]. We included those that met the pathological criteria for Alzheimer’s disease or for Dementia with Lewy bodies (DLB) categorized with neuropathological changes (NC) and who were either symptomatic or asymptomatic (with cognitive symptoms) based on the last exam prior to death, in accordance with criteria described in McKeith et al [80, 81]. Cognitive symptoms in ROSMAP included four broad categories: level 1 was assigned to those with no cognitive impairment, levels 2/3 were for those with mild cognitive impairment, levels 4-5 were for those with Alzheimer’s disease, that do not have other causes of cognitive impairment or do have other causes of cognitive impairment, respectively, and level 6 was for those with other dementias. ROSMAP participants were thus divided into five mutually exclusive categories, including Controls (not cognitively impaired), Dementia with Lewy bodies with neuropathological changes (NC) who were either symptomatic or asymptomatic (DLB-NC SYM or AD-NC ASYM, respectively), and Alzheimer’s disease with neuropathological changes (NC) who were either symptomatic or asymptomatic (AD-NC SYM or AD-NC ASYM, respectively). We additionally included combined AD and DLB traits which was a combination of their respective subgroups regardless of cognitive impairment (**Table S1**), and included amyloid beta protein levels which were measured via immunohistochemistry and quantified by image analysis and recorded as the mean amyloid beta score in 8 brain regions [82]. BMD was collected from the heel (right calcaneus) of a subset of these participants (n=114) using the Sahara Clinical Bone Sonometer at an annual visit [83].

In the Oslo study, an osteoporosis diagnosis was made according to the World Health Organization’s operational definition (bone density of ≤2.5 deviations below the mean of a 30-year-old man/woman or T score ≤2.5) [51]. BMD of the hip at the femoral neck and total body were assessed using dual-energy X-ray absorptiometry (DXA) using a Lunar Prodigy densitometer [49]. Both femoral neck T score and the dichotomous osteoporosis variable were used as separate traits in our analysis, despite the synonymity, as those who were diagnosed as osteopenic were not included under the dichotomous trait, thereby making their sample sizes different.

### Inclusion criteria

In ROSMAP, participants were required to have non-missing age, sex, postmortem interval (PMI), and RIN, resulting in a sample size of 629 for the full set. We additionally limited to individuals identified into one of the five ADRD groups described previously (**Table S1**), resulting in a sample size of 298. The smaller set of 298 was used in our primary analysis, and as an additional analysis we included the full set of unselected 629. In the Oslo study, the participants were a subset of the full study that had bone RNA array data available, resulting in a sample size of 84 (**Table 1**).

### Data preprocessing and preparation

In ROSMAP, data was log transformed and adjusted for age, sex, PMI, and RIN score with a linear model. Adjusted data was variance filtered to include genes in the top quarter (25%) of variance. In Oslo, data was log transformed and adjusted for age with a linear model. Adjusted data was also variance filtered to include genes in the top quarter (25%) of variance (**Figure 1A**).

### Construction of WGCNA networks

The WGCNA package in R (v.3.5) was used for network construction [50]. Unsigned and scale-free networks were used as the basis for both networks. A soft threshold value of 6 was selected for ROSMAP and 7 for Oslo. A similarity matrix between each pair of genes (*i* and *j*) was calculated based on the Pearson’s correlation value. The similarity matrix was transformed into an adjacency matrix by raising the correlation coefficient to the power threshold β,

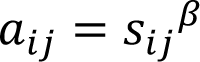

where *a_ij_* is the weighted value of the protein in the adjacency matrix defined by raising the co-expression similarity *s_ij_* to a power of *β*. Next, the topological overall matrix (TOM) and the corresponding dissimilarity (1-TOM) was calculated. Gene co-expression modules were obtained by using the dynamic cut tree method on hierarchical clustering branches [50]. Modules were not merged to retain smaller module sizes. A coloring system, innate to the WGCNA package, was used to label the modules, with modules assigned a color at random.

### Module-trait relationship analysis

Module Eigengenes (MEs), the first principal component of the module, were calculated by the WGCNA package [50] and were correlated with respective traits from the ROSMAP and Oslo studies, yielding a correlation coefficient *r* and p value *p.* To be considered a module of interest, in ROSMAP, MEs must be significant for heel BMD and at least two ADRD traits whereas in Oslo the MEs must be significant for OP and at least one other BMD measure.

Modules and their gene members from each network were overlapped to analyze correspondence of modules between the Oslo and ROSMAP networks. *p* values were calculated through a chi squared test or Fisher’s exact test and used to assess significant overlap between gene members of corresponding modules in Oslo or ROSMAP. This analysis was repeated for the full ROSMAP dataset (n=629).

### Functional enrichment analyses and module annotation

Modules from each network were analyzed with gene ontology (GO) functional enrichment analyses using the clusterProfiler (v.4.0) R package with *p*<0.05 as the significance threshold [84]. The resulting GO terms for each module were summarized on a module-level and grouped into 6 major physiological categories for the ROSMAP study and 5 major physiological categories for Oslo study in a similar fashion as a previous study [46]. This analysis was repeated for the full ROSMAP dataset (n=629), however only modules of interest were annotated.

### Signaling enrichment analysis

The term “signal” was queried in the GO pathway enrichment analysis output for modules of interest. Terms were plotted to show their respective module colors, gene counts, and *p* values, and arranged based on their levels of significance.

### Wnt term gene discovery

Pathway analysis results for ROSMAP and Oslo networks were used to extract genes that were associated with each GO term that included the term “Wnt”. Results were plotted as a tileplot to indicate important Wnt related genes in modules of interest from each network.

## Acknowledgements

We would like to acknowledge our wonderful colleague and lab member Carmen Khoo whom we lost in October 2022. Her support, hard work, and dedication to the Lary lab was pivotal to our growth and she will remain deeply missed.

We would also like to express our thanks to Meghan Gerety for her time at the lab and her help with our WGCNA pipeline development.

**Disclosures:**

