## Supplementary Figures for "Transcriptomic network analysis of brain and bone reveals shared molecular mechanisms underlying Alzheimer’s Disease and related dementias (ADRD) and Osteoporosis"

#### ROSMAP Full Group (n=629)

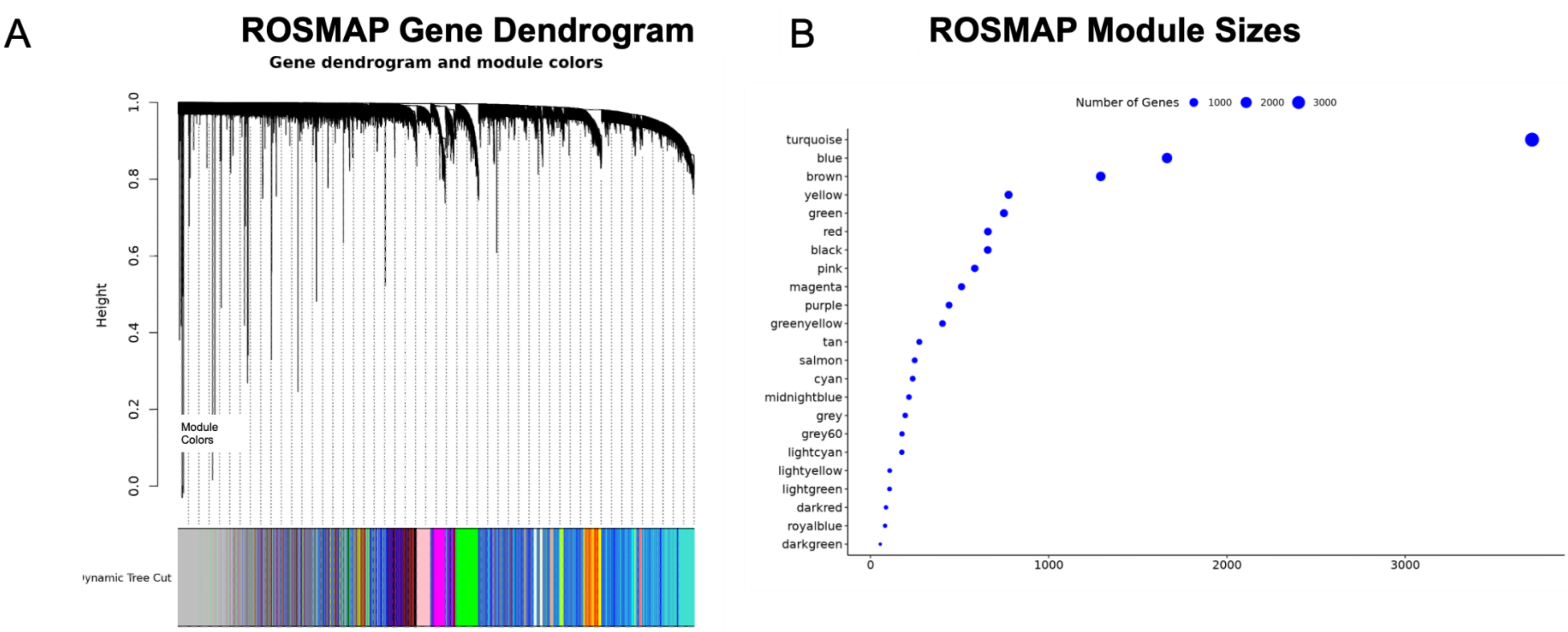


**Figure S1. WGCNA pipeline and network construction for ROSMAP (full).** (A) Gene dendrogram of ROSMAP (full) network, where colors represent the labels of modules generated in the network. (B) Plot of number of genes that correspond to each module in ROSMAP's (full) network.

### SUPPLEMENTARY FIGURES

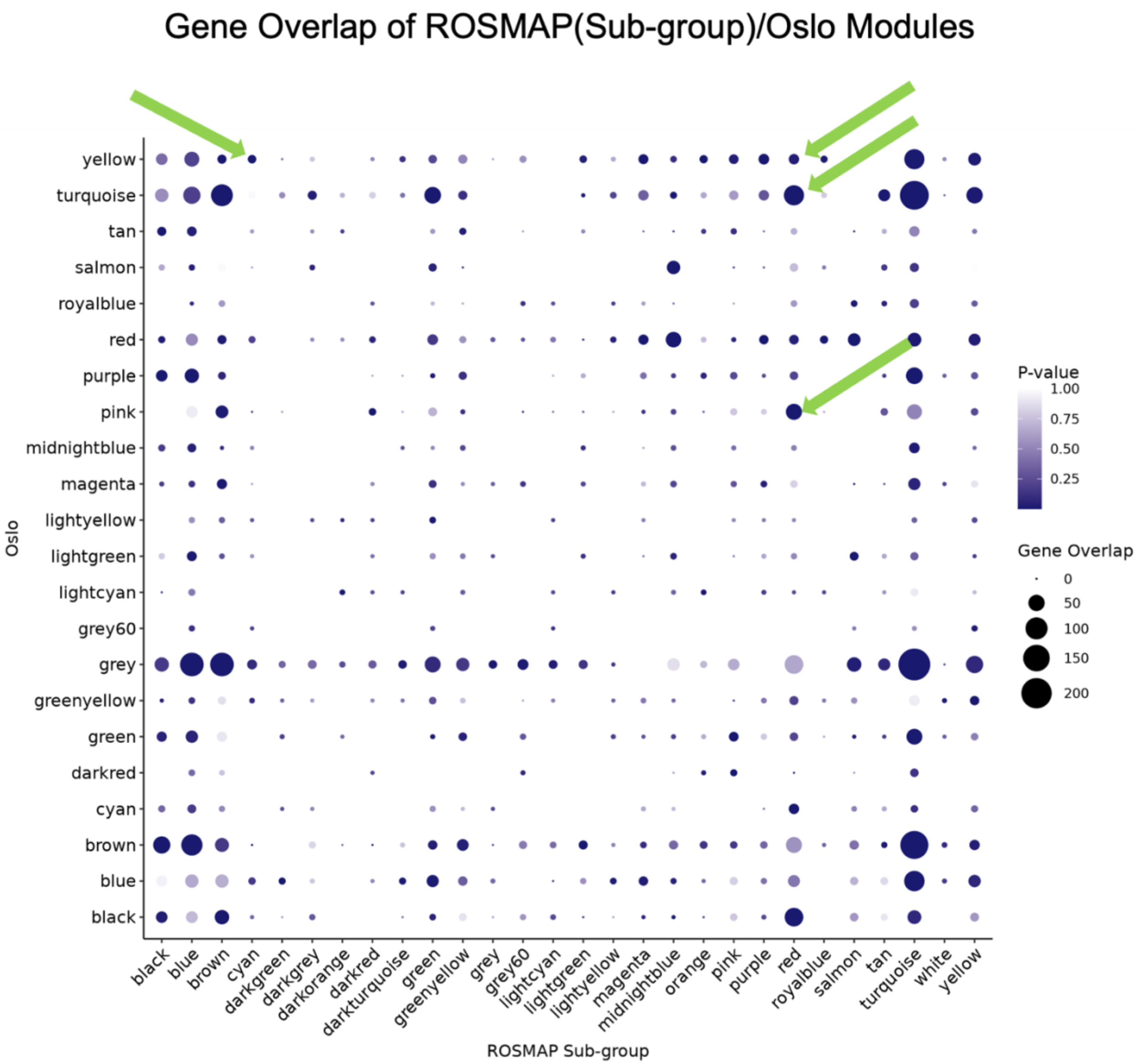


**Figure S2. Significance of gene overlap between ROSMAP (subgroup) and Oslo network modules.** Dotplot of gene overlap between Oslo and ROSMAP networks. Gene overlap is noted through dot size and *p* value through dot color**.** Green arrows highlight the overlap between modules of interest between networks.

### SUPPLEMENTARY FIGURES

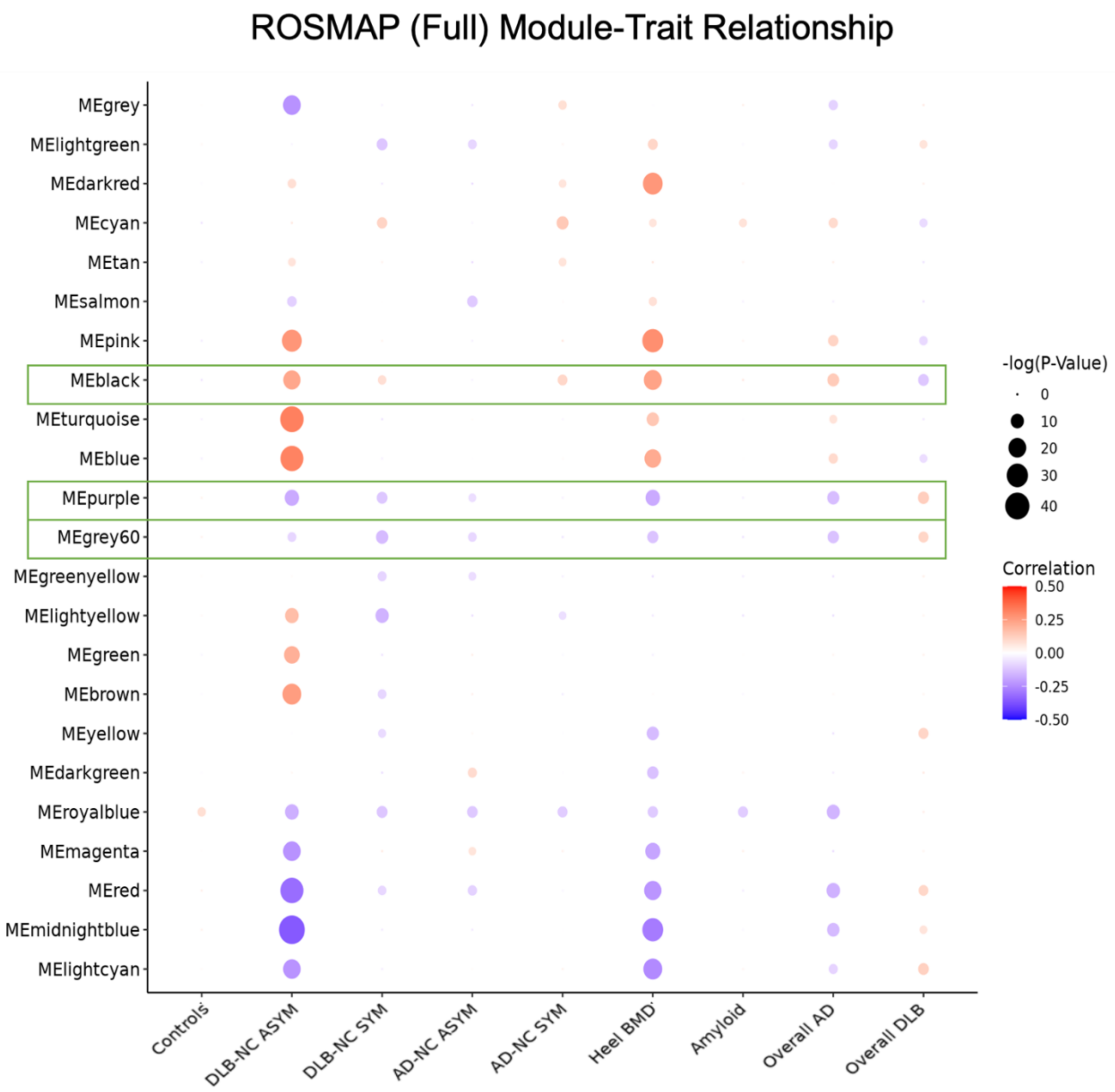


**Figure S3. Module-trait analysis of ROSMAP (full) network.** (A) Dotplot of module eigengene correlation with ADRD and bone traits in ROSMAP (full) network. Correlation values are noted through dot color and *p* value through dot size. Values shown for -log(p), where *p* < 0.1. Red boxes represent modules of interest.

### SUPPLEMENTARY FIGURES

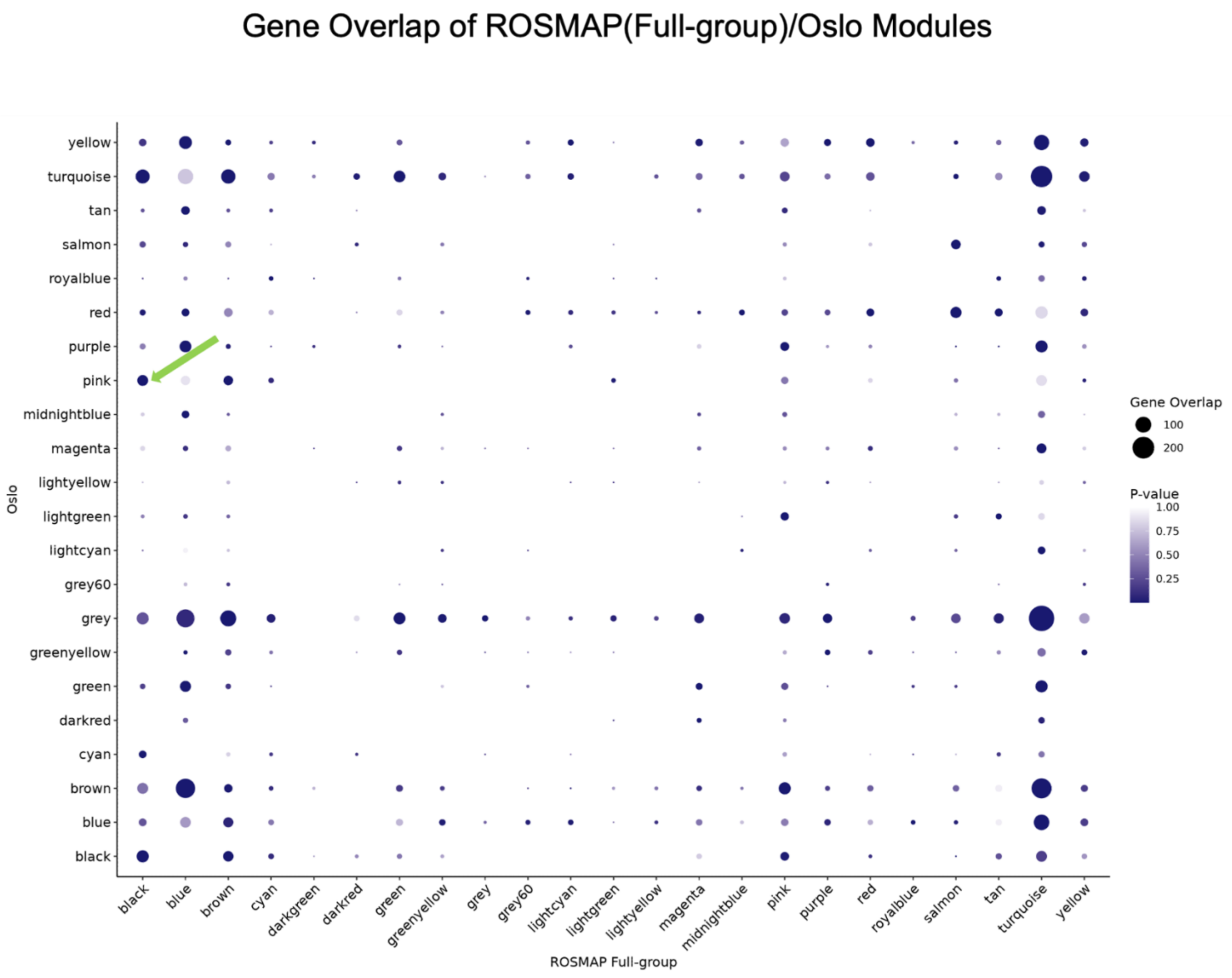


**Figure S4. Significance of gene overlap between ROSMAP (full) and Oslo network modules.** Dotplot of gene overlap between Oslo and ROSMAP (full) networks. Gene overlap is noted through dot size and *p* value through dot color**.** Green arrows highlight the overlap between modules of interest between networks.

### SUPPLEMENTARY FIGURES

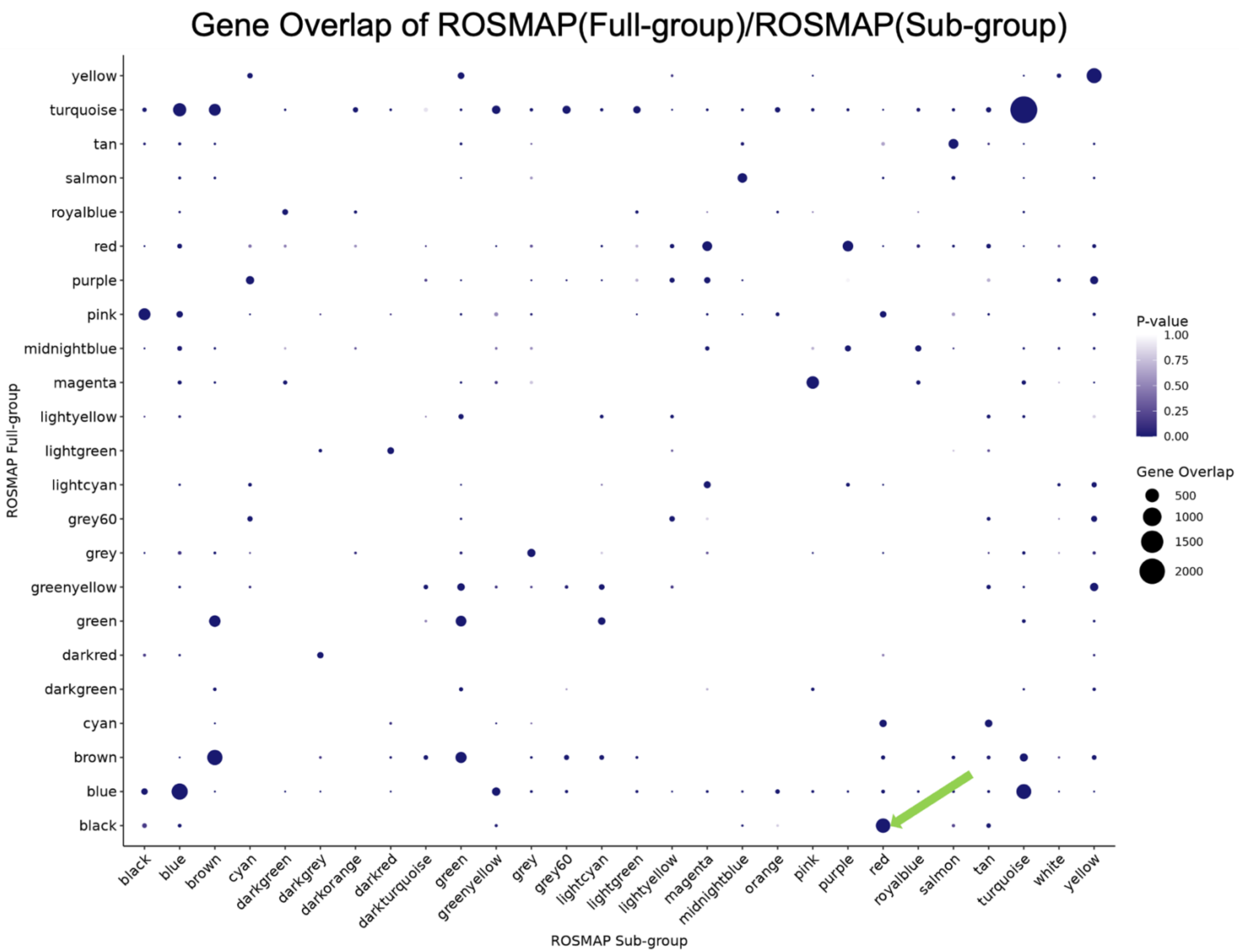


**Figure S5. Significance of gene overlap between ROSMAP (full) and ROSMAP (subgroup) network modules.** Dotplot of gene overlap between ROSMAP sub-group and ROSMAP full networks. Gene overlap is noted through dot size and *p* value through dot color**.** Green arrows highlight the overlap between modules of interest between networks.

### SUPPLEMENTARY FIGURES

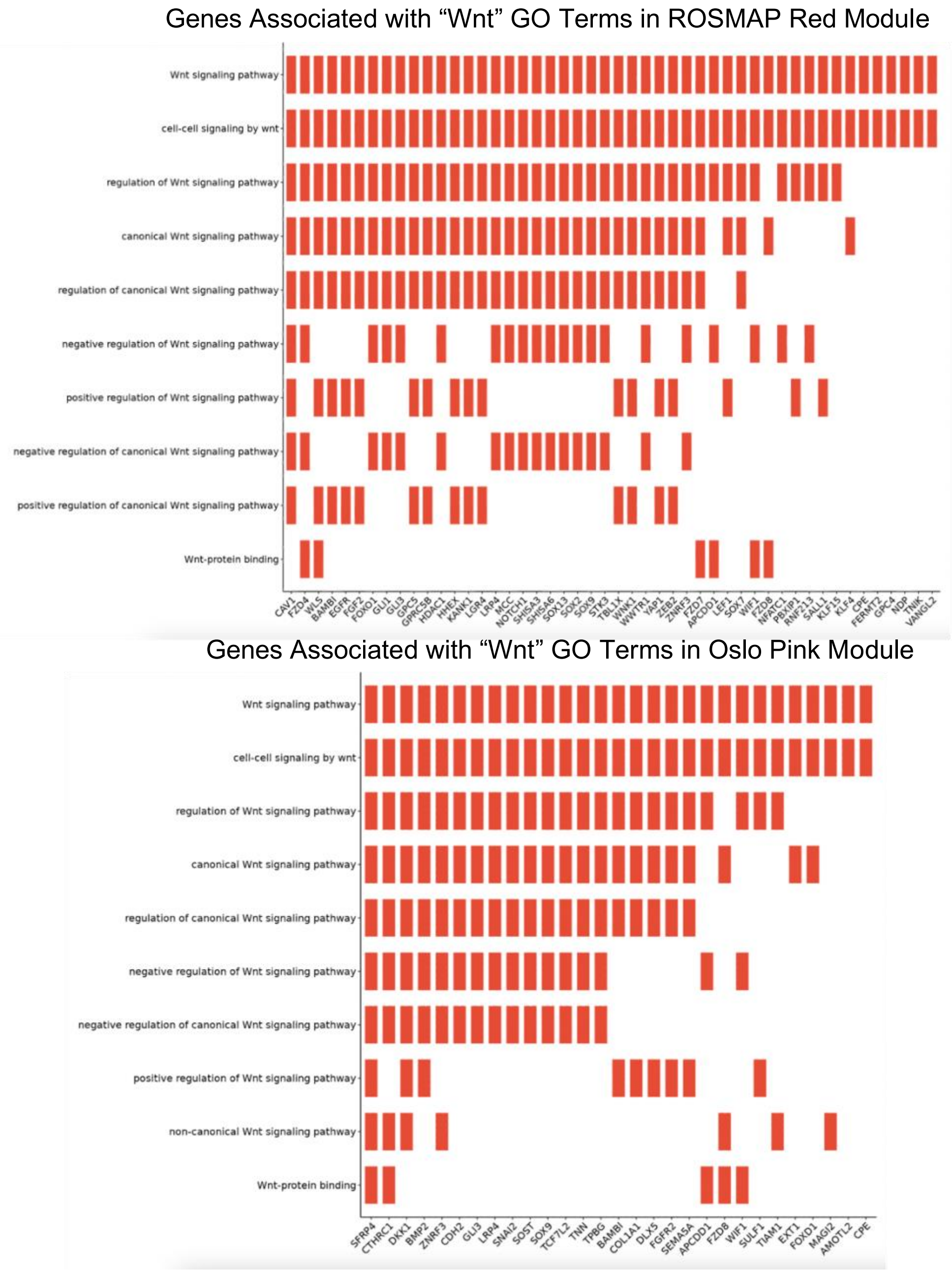


**Figure S6. Enrichment of Wnt Terms in ROSMAP red and Oslo pink modules.** (A) Tile plot of genes associated with any Wnt related GO term in the ROSMAP red module. (B) Tile plot of genes associated with any Wnt related GO term in the Oslo pink module. Green arrows highlight the overlap between modules of interest between networks.
